# Auditory Working Memory Mediates the Relationship Between Musicianship and Auditory Stream Segregation

**DOI:** 10.1101/2024.10.24.619127

**Authors:** Martha Liu, Isabelle Arseneau-Bruneau, Marcel Farrés Franch, Marie-Elise Latorre, Joshua Samuels, Emily Issa, Alexandre Payumo, Nayemur Rahman, Naíma Loureiro, Tsz Chun Matthew Leung, Karli M. Nave, Kristi M. von Handorf, Joshua D. Hoddinott, Emily B.J. Coffey, Jessica Grahn, Robert J. Zatorre

## Abstract

This study investigates the interactions between musicianship and two auditory cognitive mechanisms: auditory working memory (AWM) and stream segregation. The primary hypothesis is that AWM mediates the relationship between musical training and enhanced stream segregation capabilities. Two groups of listeners were tested, the first to establish the relationship between the two variables and the second to replicate the effect in an independent sample. Music history and behavioural data were collected from a total of 145 healthy young adults with normal binaural hearing. They performed a task that requires manipulation of tonal patterns in working memory, and the Music-in-Noise Task (MINT), which measures stream segregation abilities in a musical context. The MINT task expands measurements beyond traditional Speech-in-Noise (SIN) assessments by capturing auditory subskills (e.g., rhythm, visual, spatial, prediction) relevant to stream segregation. Our results showed that musical training is associated with enhanced AWM and MINT task performance, and that this effect is replicable across independent samples. Moreover, we found in both samples that the enhancement of stream segregation was largely mediated by AWM capacity. The results suggest that musical training and/or aptitude enhances music-in-noise perception by way of improved AWM capacity.

## Introduction

Navigating the symphony of sounds that simultaneously converge upon our ears poses a multifaceted challenge to the human auditory system. Sounds from multiple sources are combined into a single acoustic waveform, making it difficult to distinguish distinct perceptual objects. The cognitive mechanisms of auditory stream segregation involve the disentanglement of a target auditory signal from a web of ambiguous and irrelevant background noises (Bregman, 1990), thus playing a pivotal role in organizing our auditory input (Shamma & Micheyl, 2010). This advanced cognitive function is influenced by both stimulus-driven and cognitive top-down factors (Anderson et al., 2013; Davis & Johnsrude, 2007). Consequently, unravelling the nuances in stream segregation carries important implications for understanding auditory cognitive processes and refining auditory neurocognitive models.

At the sensory level, simultaneous and sequential grouping strategies (e.g., timbre similarity) constitute the basic perceptual aspects of auditory stream segregation (Noorden, 1975; Deroche et al., 2017; Bregman & Pinker, 1978). At the cognitive level, stream segregation involves various factors such as the listener’s attention and attentional load (Heinrich et al., 2008; Thompson et al., 2011), working memory skills (Escobar et al. 2020), prior knowledge (Davis & Johnsrude, 2007), and schematic expectations (Bey & McAdams, 2002). In particular, auditory working memory (AWM) has been suggested to play a crucial role in auditory stream segregation and accounts for individual differences in this capacity (Gordon-Salan & Cole, 2016; Parbery-Clark et al., 2009a).

### AWM and Stream Segregation

Evidence for the involvement of AWM in stream segregation stems from studies showing the influence of predictive mechanisms on this ability. Bey and McAdams (2002) demonstrated that stream segregation is improved when a target melody is heard in advance of the stimulus presented in noise, suggesting that holding the target in short-term memory facilitates its subsequent anticipation and detection. In addition, the role of AWM in stream segregation is supported by studies showing that working memory modulates speech perception and attention in noisy environments (Heinrich et al., 2008). Heinrich et al. found that extracting speech from noise places demands on the listener’s attentional capacity, resulting in a decrease in the availability of other mental processing. However, enhanced AWM proficiency facilitates the encoding of the acoustic signal, thus allowing greater allocation of cognitive resources toward extracting information and recalling target words. Furthermore, Dalton et al. (2009) presented evidence supporting the causal role of working memory in diminishing interference in an auditory task involving distractor stimuli.

### The Influence of Musicianship on AWM and SIN Perception

Musicians have emerged as a distinctive population of interest in AWM and stream segregation research due to their constant exposure to and attunement to complex auditory streams (Brown et al., 2015). On one hand, their capacity to discern and organize auditory signals is finely tuned in musical performance, leading to improved stream segregation abilities (Swaminathan et al., 2015; for review, see Coffey et al. 2017b). On the other hand, musicians extensively engage their AWM during the process of learning and practicing, such as retaining tunes and rehearsing melodies. A meta-analysis shows that musicians outperform non-musicians in working memory, especially when the stimuli are tonal (Talamini et al., 2017). Studies have consistently demonstrated behavioural and neurophysiological evidence for the positive influence of music experience on standardized measures of AWM (for review, see Yurgil et al., 2020). Such observed advantages among musicians make them well-suited for the examination of the interaction between musicianship, AWM and stream segregation.

Traditionally, the relationship between these two variables has been examined using a variety of Speech-in-Noise (SIN) tests that involve hearing target speech phrases or words at different signal-to-noise ratios (SNRs; Nilsson, 1994; Killion et al., 2004). With the SIN measure, a large number of studies have reported that musical training is correlated with a better perception of speech-in-noise (for review, see Coffey et al. 2017b) and is associated with differences in AWM (Kraus et al., 2012). Research by Parbery-Clark et al. (2009b) showed that both younger and older musicians outperform age-matched non-musicians on the standardized measures of SIN perception. In addition, musicians in that study were shown to have better performance on an AWM task involving the recall and reordering of a series of numbers and words. Based on the analysis of the strong relationship between AWM and SIN perceptual abilities across age groups, the researchers suggested that the AWM enhancement of musicians mediates their better performance in SIN (Parbery-Clark et al., 2009b).

Despite the convincing evidence and theory, musician enhancements on either AWM or SIN have not been consistently reported. Some studies show no difference between musicians and non-musicians in AWM ability (Escobar et al., 2020), while others demonstrate similar insignificant findings for SIN perception (Boebinger et al., 2015; Madsen et al., 2019). A study by Escobar et al. (2020) indicated no significant interaction between musical training and WM, and no statistically significant advantages for musicians’ SIN perception compared to non-musicians when matched to WM ability. Research adopting Mandarin nonsense sentence stimuli has shown a mediating role of AWM in ameliorating SIN perception loss in older musicians, while such musician advantages were absent in young adults (Zhang et al., 2021). Other research reported musicians’ SIN advantage and correlation between SIN scores and working memory, although the associations are limited to cases where the noise induces linguistic interference (Yoo & Bidelman, 2019).

The contentious findings in SIN tests could be related to variations in task design, criteria for musicianship, and different scoring methods. More critically, these SIN tasks fall short of providing the granularities required to assess individual perceptual components and top-down cues involved in stream segregation, which could potentially be affected by training or other interventions. Furthermore, SIN assessments in prior studies exclusively focused on sentence or word detection, which were related to linguistic skills and schematic knowledge (for a review, see Coffey et al., 2017b). This focus potentially limits the generalizability of the findings on hearing-in-noise to the speech modality alone.

### Music-in-Noise Task (MINT)

To eliminate linguistic influences and evaluate different auditory sub-skills in stream segregation, Coffey et al. (2019) introduced the Music-in-noise task (MINT). MINT expands measures beyond speech perception by incorporating a melodic line as the target, enabling the examination of selective listening of musical phrases amidst a mix of musical sounds that provides informational masking. By employing simple melodies as the target signal, MINT allows for the disentanglement of the relative contributions of top-down processes and the systematic integration of rhythmic, visual, spatial attentional and predictive cues, which have been shown to be crucial in stream segregation (Slater & Kraus, 2016; Coffey et al., 2019). Paralleling the findings in SIN research, Coffey et al.’s study using the MINT task revealed significant correlations between cumulative hours of musical practice and music-in-noise perception, particularly in rhythm, prediction, and visual conditions. The study also showed a significant relationship between AWM and overall MINT performance.

Nevertheless, in Coffey et al.’s (2019) study, AWM capability was only accounted for as a covariate in analyzing musical training’s impact on MINT sub-conditions, along with other factors such as pitch perception and multilingualism. Consequently, there remains a gap in the literature regarding the interaction between musical training, AWM, and music-in-noise perception. Here, we aim to shed light on the underlying dynamics of how musical experience influences auditory perceptual and memory processes.

### Specific Aims and Hypothesis

The goal of the present study was to determine if the purported musician advantage in auditory stream segregation could be consistently observed, and specifically to test the hypothesis that such an effect is mediated by enhanced AWM. We implemented a test-replication research design where the same study is conducted in two phases with independent samples. This approach allows for testing the robustness of the findings across cohorts of different distributions of musicianship. In Experiment 1 (Initial Phase) the phenomenon of interest is identified and analyzed. while Experiment 2 (Replication Phase) tests whether the initial findings can be replicated, thus ensuring that the observed effects are robust and not solely related to the specific sample used in the first phase.

To test the effects of music training on both MINT and AWM we carried out correlational analyses using years of musical training as the independent variable; for additional verification and to account for possible nonlinear effects we also carried out categorical comparisons of musicians vs nonmusicians. We hypothesized a positive relationship between musical training and AWM and MINT task performance. Finally, we aimed to test the hypothesis that musical training fosters improvements in MINT through the enhancement of AWM capabilities, as suggested but not fully confirmed by the literature, positioning AWM as a mediating factor in this relationship. We therefore used statistical mediation analysis, to understand the underlying process by which musical training influences music-in-noise perception, delineating direct and indirect effects through the mediator (AWM).

## Experiment (1)

### Methods and Materials

#### Participants

In the initial phase, we recruited 82 healthy young adults (30 males, 51 females, 1 non-binary) with either minimal or extensive piano experience (total practice hours: mean = 5300, *SD* = 5900, range: 0-30000). The average age was 25.5 years (*SD* = 6.8, range: 18-45). To conduct group comparisons on the effects of musical training, we defined subjects with >10 cumulative years of music training and > 4000 h of lifetime practice as Musicians (*N* = 42; mean practice hours: 9400, *SD* = 5700), and subjects with <2 years of musical activity as Non-Musicians (*N* = 20, mean practice hours: 94, *SD* = 203). For the Musicians, the average age of onset for music training is 5.0 (*SD* = 1.3, range: 2-7).

Subjects provided informed consent and were compensated for their participation and time. All experimental procedures were approved by the McGill University Faculty of Medicine Research Ethics Board. All participants were screened to have normal or corrected-to-normal vision and reported no history of neurological disorders. Normal binaural hearing was confirmed by an audiometric test which measured pure-tone thresholds from 250 to 8000 Hz (less than 25 dB SL pure-tone hearing thresholds between 125 and 8000 Hz). Participants with binaural hearing thresholds above 25 dB SL did not proceed with the study as deficiencies in the frequency range may influence their task performance. Out of the 82 participants from Experiment 1 who completed all parts of the study, 4 were excluded from the MINT analysis due to their inability to process basic musical content (with 2 or more out of 6 incorrect responses for the MINT task Control condition, see description below).

#### Procedure

Prior to the testing session, participants confirmed eligibility and completed the Montreal Music History Questionnaire (MMHQ; Coffey et al., 2011). The MMHQ provides the subject’s self-reported information regarding overall musical experience (instruments played, total cumulative practice hours), language proficiency, basic demographics, etc. The testing session began with an audiometry hearing test, followed by the two behavioural tasks: the AWM task (Albouy et al., 2017), and the MINT task (Coffey, 2019); see the following section for descriptions. The visual component of each task was presented on a computer screen and sounds were presented binaurally through headphones (ATH-M50x, Audio-Technica). A comfortable sound level was determined during pilot testing and kept constant for all subjects and both tasks.

#### Measures and Behavioural Tasks

(1) To test for individual AWM abilities and eliminate linguistic influences, we implemented an AWM task that measures individuals’ auditory retention and manipulation capabilities with sets of tonal stimuli (Albouy et al., 2017). This AWM task uses a discrimination design that involves the detection of a local pitch change within two tonal patterns differing in temporal order, described as the ‘Manipulation Task’ in Albouy et al. (2017). On each trial, participants first listened to three sequentially presented 250 ms tones, which were followed after a 2000 ms silent retention interval by a probe consisting of another set of three tones (**Figure 1**). The task was to determine whether the sequence of the second set of three tones was a perfect reverse of the first set or not. The structure of this task engages AWM capabilities, requiring participants to retain the initial set of tones and inversely manipulate them in their mental workspace during the retention interval (Albouy et al., 2017; Foster et al., 2013; Zatorre et al., 2010). 6 practice trials with feedback were provided, followed by 100 experimental trials without feedback. Task trials are randomized with a maximum of 3 consecutive trials with the same condition. The average accuracy score was then computed based on the percentage of responses correct.
(2) The Music-in-Noise Task (MINT) assesses stream segregation, involving the detection of a target musical melody embedded in irrelevant musical background noise (Coffey et al., 2019). Employing a match-mismatch discrimination design, each trial features one melodic line embedded in masking noise, and a melodic line presented in silence (**Figure 2**). Participants were asked to judge if the two presented melodies were the same or different. The MINT consists of five conditions which capture auditory sub-skills and the influence of perceptual cues: (1) Baseline (Pitch; **Figure 2A**), where the target-noise mixture is first presented, followed by the comparison melody in silence, without additional cues; (2) Rhythm (**Figure 2B**), the target is a rhythmic pattern with no pitch variation; (3) Spatial (**Figure 2D**), an additional spatial attentional cue is presented for the participant to attend to sounds coming from their left or right side (the perception of which is manipulated via interaural sound level difference); (4) Visual (**Figure 2E**), an additional visual cue outlining the melody’s contour is presented to facilitate target detection within the mixture; and (5) Prediction (**Figure 2C**), subjects hear the target melody in silence first, followed by the comparison melody in noise. There is also a control condition with both melodies presented in silence to screen out participants incapable of discriminating the musical content of the MINT task, and who may therefore have amusia (Peretz et al. 2002). All conditions were tested at three different signal-to-noise (SNR) levels (0, −3, and −6 dB). Each condition involved 2 practice trials, followed by 20 experimental trials presented in a randomized block order across subjects. The accuracy score for each individual condition and overall performance is calculated by averaging the percentage of correct responses across all SNR levels within the respective condition(s); and the accuracy score for performance at each SNR level is computed by averaging the percentage of correct responses across all conditions at that specific SNR level (for further procedural details, see Coffey et al., 2019).

## Results

### Musical Training and AWM

The mean accuracy score (% correct) for the AWM task was 78.8 (*SD* = 18.5, *N=78*). Pearson correlation indicates a significant relationship between cumulative hours of practice and AWM task performance (*r* = 0.399, p < 0.001; **Figure 3**). An independent samples t-test was conducted to examine the group difference between Musicians and Non-Musicians on AWM score. There was a significant difference in AWM abilities between Musicians (mean = 86.94, *SD* =13.11) and Non-Musicians (mean = 56.65, *SD* = 12.57); t(60) = –8.62, p < .001 (**Figure 5**).

**Figure 1.**
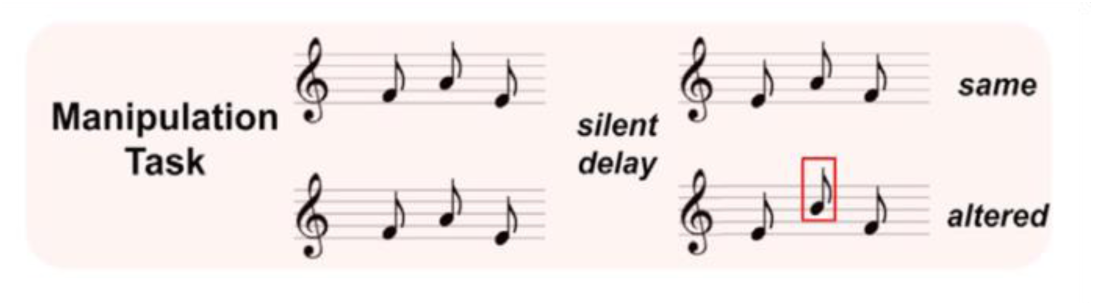
Illustration of AWM task (Adapted from Albouy et al., 2017). ‘Match’ trials: the second sequence of melody was presented in a reversed temporal order of the first melody; ‘mismatch’ trials: second melody was presented in reversed temporal order, with one local pitch change. This required the retention and manipulation of auditory information.

**Figure 2.**
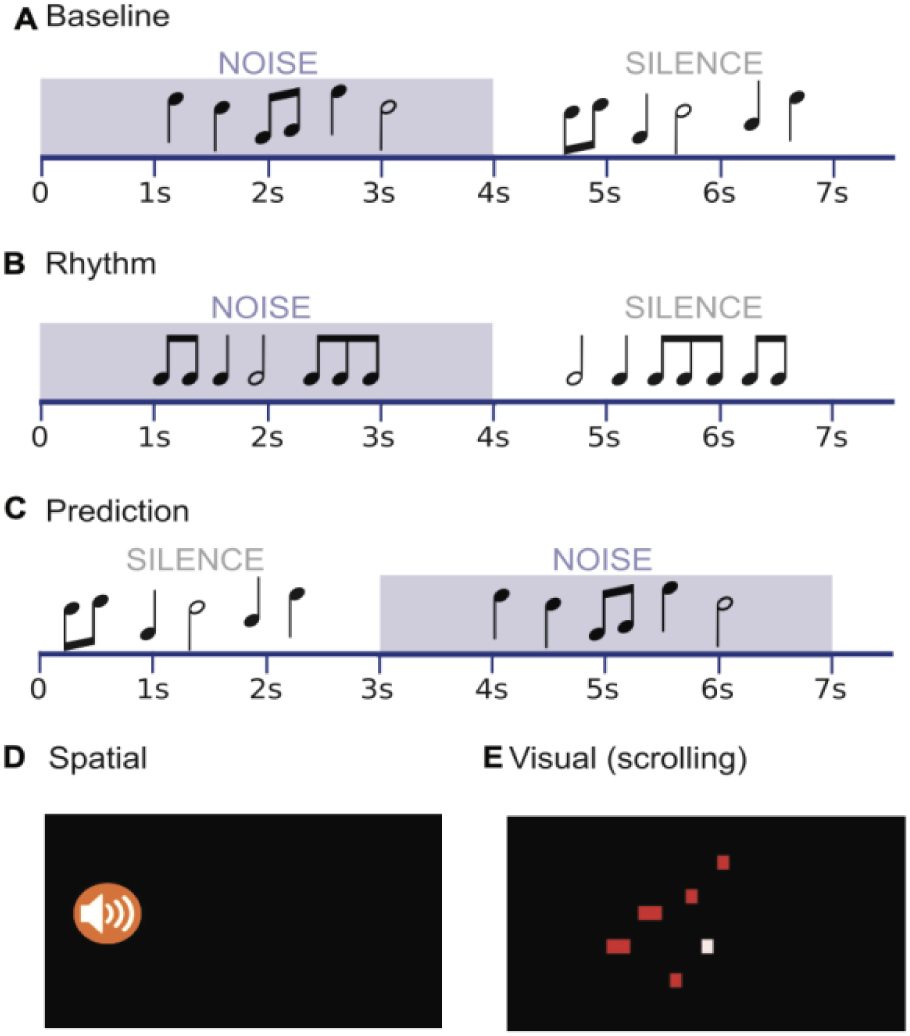
Illustration for MINT (Adapted from Coffey et al., 2019). ‘Match’ trials: the melody mixed in noise is identical to the melody presented in silence; ‘mismatch’ trials: the melody mixed in noise is not identical to the melody presented in silence. MINT consists of five conditions: (A) Baseline (Pitch), (B) Rhythm, (C) Prediction, (D) Spatial, and (E) Visual. In the Spatial condition (D), an icon on one side of the screen directed the listener to attend to the corresponding ear. In the Visual condition (E), a scrolling graphic representation outlines the timing and melodic contour of the target melody.

**Figure 3.**
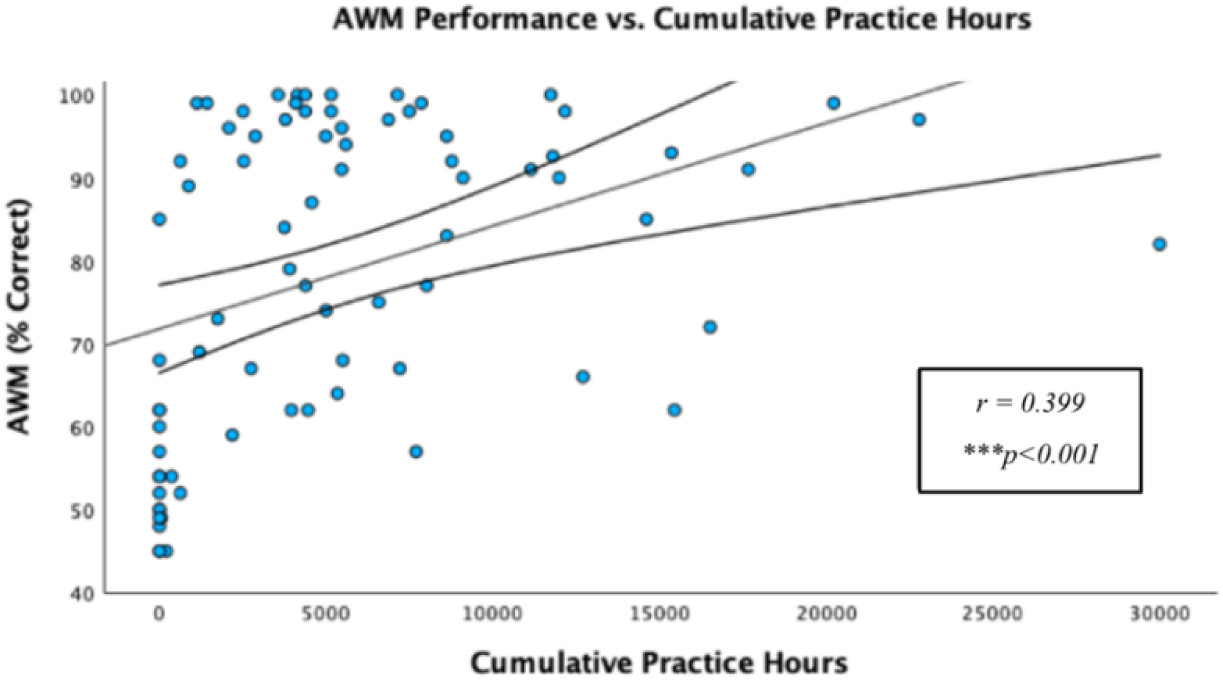
Experiment 1 cumulative practice hours vs. AWM task performance. Pearson correlation is significant at the 0.01 level).

### Musical Training and MINT Outcomes

The mean accuracy score (% correct) for the overall MINT was 82.05 (*SD* = 760). The mean accuracy scores for each MINT sub-condition were: Baseline (Pitch) = 80.94 (*SD* = 11.95), Rhythm = 63.85 (*SD* = 14.56), Spatial = 84.02 (*SD* = 10.49), Visual = 90.60 (*SD* = 10.27), and Prediction = 90.85 (*SD* = 8.80). The mean accuracy scores for each SNR level were: SNR 0 = 84.77 (*SD* = 10.74), SNR –3 = 83.79 (*SD* = 9.32), and SNR –6 = 77.59 (*SD* = 9.93). Pearson correlation analysis between cumulative practice hours and overall MINT task performance revealed a significant correlation, with a *r* value of 0.363 (p < 0.001; **Figure 4**). Cumulative hours of practice were also correlated with the Baseline (Pitch) (*r* = 0.22, p = 0.025), Prediction (*r* = 0.26, p = 0.010), Rhythm (*r* = 0.28, p = 0.007), and Visual (*r* = 0.29, p = 0.005) sub-conditions. In addition, cumulative hours of practice correlated with all SNR levels: SNR 0 (*r* = 0.24, p = 0.019), SNR –3 (*r* = 0.31, p = 0.003), and SNR –6 (*r* = 0.29, p = 0.006). Independent samples t-test shows a significant difference in MINT performance between Musicians (mean = 84.63, *SD* =5.86) and Non-Musicians (mean = 75.13, *SD* = 8.36); t(60) = –5.18, p < .001 (**Figure 5**).

**Figure 4.**
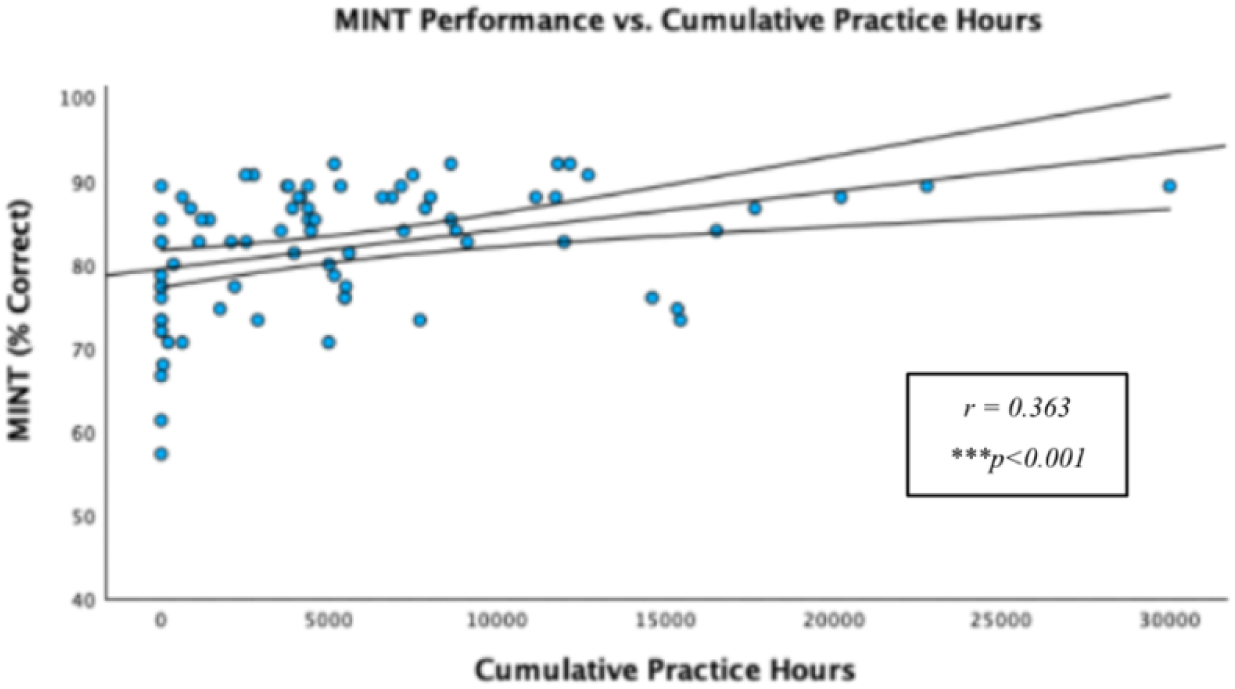
Experiment 1 cumulative practice hours vs. overall MINT performance. Pearson correlation is significant at the 0.01 level.

**Figure 5.**
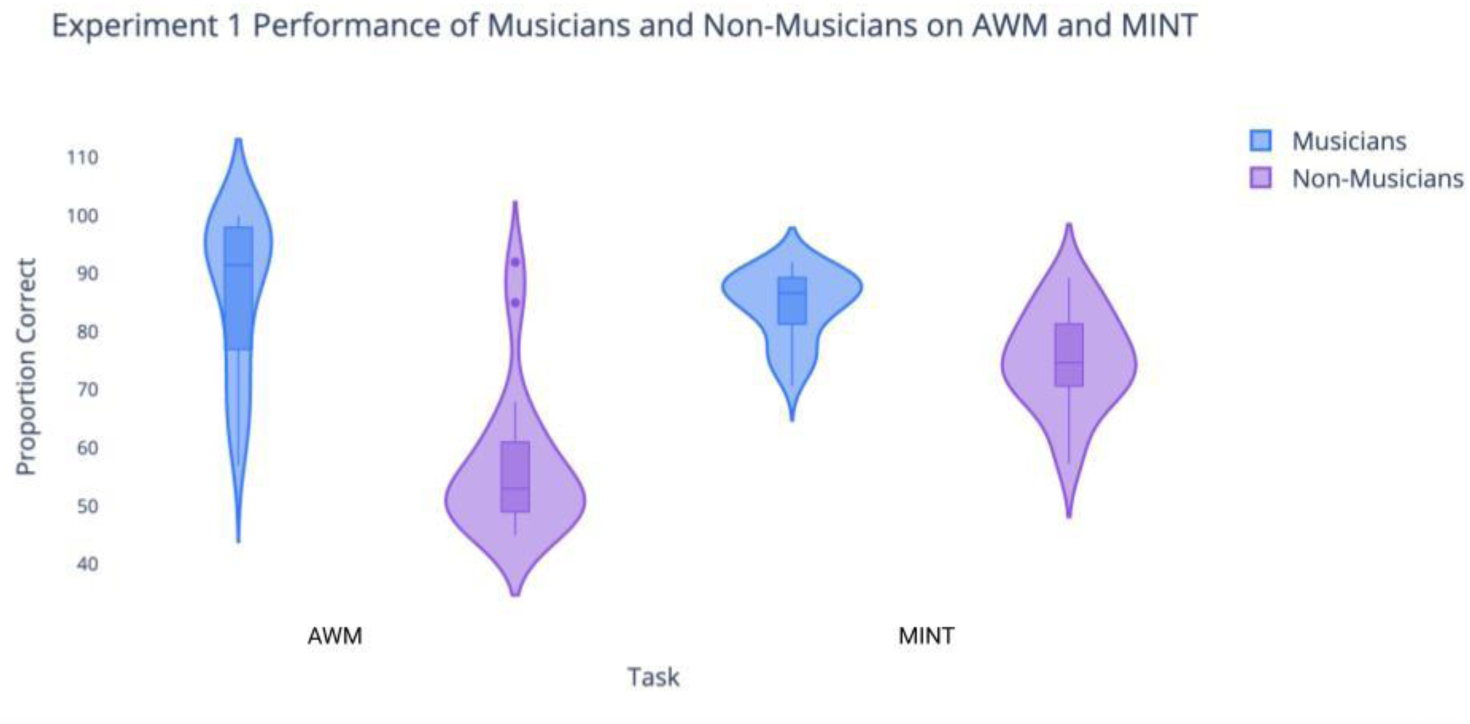
Violin plot showing Experiment 1 AWM task performance for Musician (mean = 86.94, *SD* =13.11, *N* = 42) and Non-Musician groups (mean = 56.65, *SD* = 12.57, *N* = 20). MINT performance for Musician (mean = 84.63, *SD* =5.86) and Non-Musician groups (mean = 75.13, *SD* = 8.36). Significant group difference for both tasks *p* < 0.001.

### AWM and MINT Performance

Pearson correlation analysis evaluated the relationship between performance on the AWM and MINT tasks. The AWM scores significantly correlated with the overall MINT scores (*r* = 0.584, p < 0.001; **Figure 6**). The AWM correlated with all MINT sub-conditions, as listed in **Table 1**. Moreover, AWM was correlated with all the SNR levels, as presented in **Table 2**. Fisher’s test performed to compare the differences between the z-transformations of each pair of correlations demonstrated that none of the correlations were significantly larger than the others.

**Figure 6.**
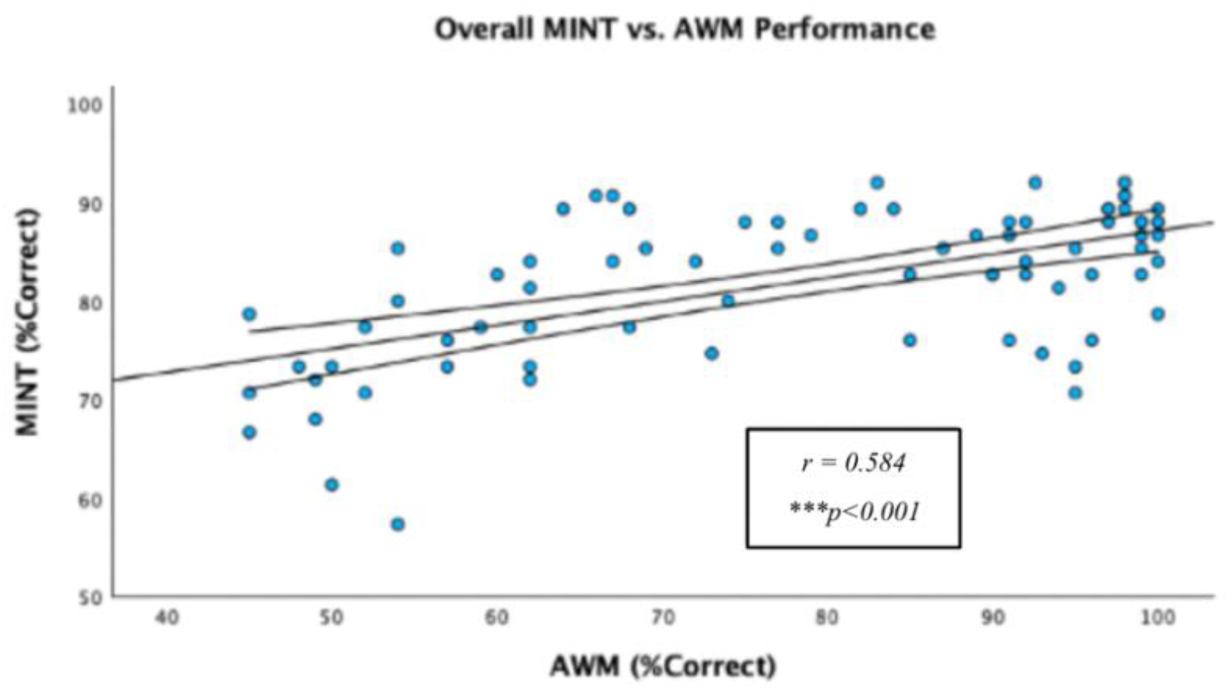
Experiment 1 AWM ability vs. overall MINT performance. Pearson correlation is significant at the 0.01 level.

**Table 1.** Experiment 1 AWM Task Performance vs. MINT Conditions. Pearson correlations based on AWM task percent correct and average MINT scores for the corresponding condition.

|  | **Pitch**<br>**Total** | **Prediction**<br>**Total** | **Rhythm**<br>**Total** | **Spatial**<br>**Total** | **Visual**<br>**Total** | **MINT**<br>**Overall** |
| --- | --- | --- | --- | --- | --- | --- |
| **AWM Task Performance** | .424** | .447** | .373** | .244** | .509** | .584** |
| **Sig. (2-tailed)** | <.001 | <.001 | <.001 | 0.032 | <.001 | <.001 |

**Table 2.** Experiment 1 AWM Task Performance vs. MINT SNR Levels. Pearson correlations based on AWM task percent correct and average MINT scores for each SNR level across conditions.

|  | **SNR-6** | **SNR-3** | **SNR 0** |
| --- | --- | --- | --- |
| **AWM Task Performance** | .538** | .360** | .432** |
| **Sig. (2-tailed)** | <.001 | <.001 | <.001 |

### Mediating Role of AWM

Regression analyses were performed to assess each component of the proposed mediation model. First, it was found that cumulative music training hours were positively associated with both MINT performance (R = 0.36, *F* (1,76) = 11.56, p = 0.001) and AWM performance (R = 0.40, *F* (1,76) = 14.42, p < 0.001). It was also found that the mediator, AWM ability, was positively related to the MINT test score (R = 0.58, *F* (1,76) = 39.42, p < 0.001). Lastly, multiple regression analysis was conducted to examine the effects of hours of musical training (*X_1_*) and AWM (*X_2_*) on MINT performance (*Y*). Results indicated that the overall regression model was significant, (R = 0.601, *F* (2,75) = 21.24, p < 0.001), with both predictors contributing to better MINT performance (β*_1_* = 0.16, *p* = 0.129; β*_2_* = 0.52, p < 0.001; **Figure 7**).

**Figure 7.**
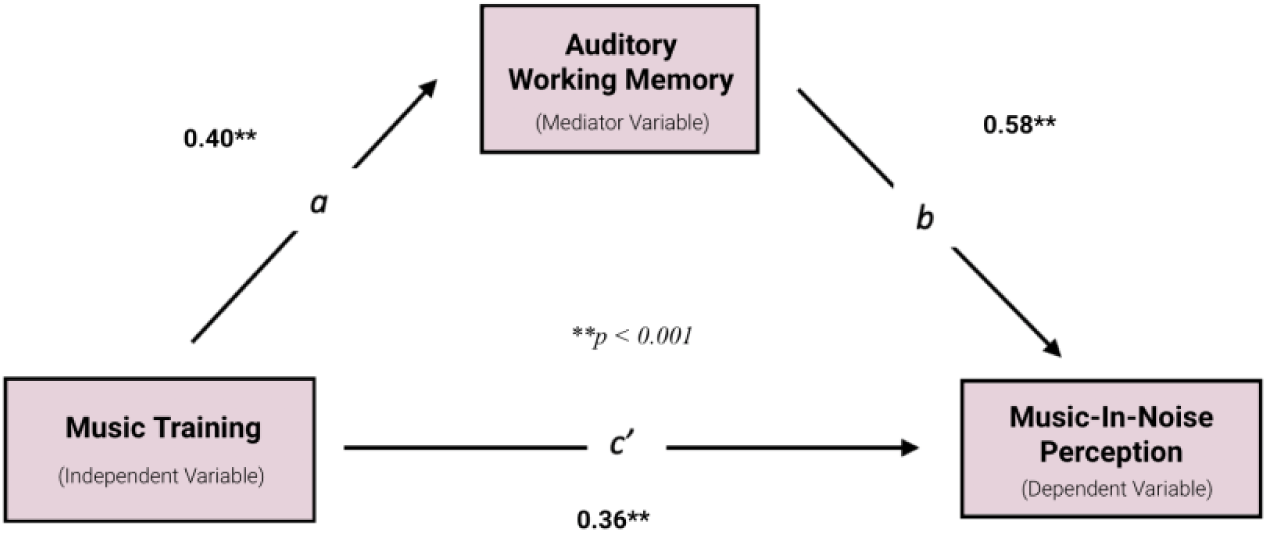
Mediation analysis results. Enhanced AWM was the significant mediator of the correlation between music training (cumulative practice hours) and MINT performance.

Because the general model, the a-path (music training to AWM), and the b-path (AWM to MINT) were significant, mediation analysis was tested using the bootstrapping method with bias-corrected confidence estimates (Preacher & Hayes, 2004). The 95% confidence interval of the indirect effect was obtained with 5000 bootstrap samples (Preacher & Hayes, 2008), and confirmed the significant mediating role of AWM in the relationship between music training and MINT task performance (**Figure 7**). Regression results also indicated that the direct effect of music training on MINT becomes non-significant (p = 0.11) when controlling for AWM, thus suggesting full mediation. Moreover, confidence intervals derived from bootstrapping mediation analysis revealed mediation effects of AWM on the Baseline (Pitch), Prediction, Rhythm, and Visual sub-conditions. Our results also indicate significant mediating effects of AWM on MINT performance at the SNR 0 and SNR –6 levels.

## Interim Discussion

The main findings from Experiment 1 aligned with our hypothesis, highlighting a clear advantage for musicians in both AWM abilities and music-in-noise perception. The results reveal a positive correlation between the number of practice hours and AWM task performance, and that the Musician group consistently outperformed Non-Musicians in AWM abilities. Additionally, both correlational and group comparison analyses illustrate a significant association between musical experience and enhanced music-in-noise performance. The bootstrapping analysis concerning practice hours, AWM and MINT further supports our mediation hypothesis, suggesting that AWM ability substantially mediates the relationship between musical experience and music-in-noise perception.

It is important to note that the majority of subjects from Experiment 1 were selected to fall into either non-musicians or expert musicians categories. Consequently, the dataset includes fewer subjects with moderate exposure to music and thus may be less reflective of the general population’s musical experience distribution. Although Pearson’s correlations indicate a notable parametric association between music training and both AWM and music-in-noise abilities, replicating the main effects observed in Experiment 1 based on a more normative and representative dataset would strengthen the statistical robustness and generalizability of the results.

In addition, results from the MINT task in Experiment 1 showed that participants performed optimally around the 80% mark, suggesting that the SNR range tested (0, –3, and –6) may not fully challenge their music-in-noise capabilities. In light of these findings, we devised a second phase of the study to extend the difficulty of the MINT task with SNR levels of –3, –6, and –9. By adjusting the noise ratio, we aim to better understand how musicianship affects MINT performance under more demanding conditions and to assess whether the effects observed in Experiment 1 persist with increased task demand. This modification should provide an assessment of the consistency of musical training effects across a wider range of noise interference challenges.

Based on the main correlational results from Experiment 1, we determined the minimum sample size required for Experiment 2 to achieve the desired statistical power. Using an expected correlation coefficient (ρ) of 0.40, a significance level (α) of 0.05, and a power (1 – β) of 0.90, and applying the Fisher Transformation of the correlation coefficient, the minimum sample size required for Experiment 2 is calculated to be 66.

## Experiment (2)

### Methods and Materials

#### Participants

In the replication phase, we recruited 73 subjects (35 males, 34 females, 1 non-binary) with a distributed range of music experience and expertise (total practice hours: mean = 4600, *SD* = 6600, range: 0-34000). Subjects from Experiment 2 have on average fewer practice hours than those in Experiment 1; t(143) = –1.84, p = 0.034. The average age was 27.0 years (*SD* = 6.6, range: 18-49). Within the 73 subjects, 19 were categorized as Musicians according to the same criteria as above (mean practice hours: 12000, *SD* = 8600), and 18 were Non-Musicians (mean practice hours: 200, *SD* = 600). For the Musicians, the average age of onset for music training is 6.0 (*SD* = 2.5, range: 2-12).

All procedures and screening criteria remained consistent with those in Experiment 1 and were approved by either the McGill University Faculty of Medicine Research Ethics Board or Western University Nomedical Research Ethics Board. Out of the 73 subjects who completed all components of Experiment 2, 3 who could not process basic musical content were excluded from the MINT analysis.

#### Procedure

Refer to Experiment 1 Methods and Materials section Procedure.

#### Measures and Behavioural Tasks

Refer to Experiment 1 Methods and Materials section Measures and Behavioural Tasks.

## Results

### Musical Training and AWM

The mean accuracy score (% correct) for the AWM task in the second sample was 66.33 (*SD* = 15.85, range: 41-100, *N* = 70). Results from a one-tailed Pearson correlation test indicated a trend toward significance in the association between musical training and AWM task performance (r = 0.191, p = 0.057). Potential outlier effects were suspected through examination of the data distribution, prompting the use of Spearman’s rank-order correlation, which is more robust to extreme values. The Spearman’s test revealed a significant monotonic relationship between AWM scores and cumulative hours of practice (ρ = 0.324, p = 0.003). The discrepancy between the rank-order and parametric test results suggests that the data may have been affected by extreme values. Upon comprehensive examination of the total 148 qualified subjects from Experiment 1 and 2 using linear regression (practice hours versus AWM performance), we identified two subjects from Experiment 2 with performance significantly deviating from the model’s predictions. Specifically, one subject had a standardized residual of –2.45 and the other – 2.40, while the standardized residuals for the remaining 146 subjects ranged between –1.67 and 1.70. Consequently, these two subjects are considered outliers and were excluded from subsequent analysis.

By removing the two outliers, the adjusted mean AWM accuracy score in the second sample task was 66.78 (*SD* = 15.83, *N* = 68). Independent samples t-test indicates a significantly lower AWM performance for the subjects in Experiment 2 compared to Experiment 1; t(144) = – 4.17, p < .001. A significant relationship between AWM score and cumulative hours of practice is demonstrated with Pearson’s test (*r* = 0.370, p < 0.001; **Figure 8**). In addition, the independent samples t-test also indicates a group difference in AWM performance between Musicians (mean = 76.68, *SD* =16.60) and Non-Musicians (mean = 55.44, *SD* = 6.82, *N* = 18); t(35) = 5.04, p < .001 (**Figure 10**).

**Figure 8.**
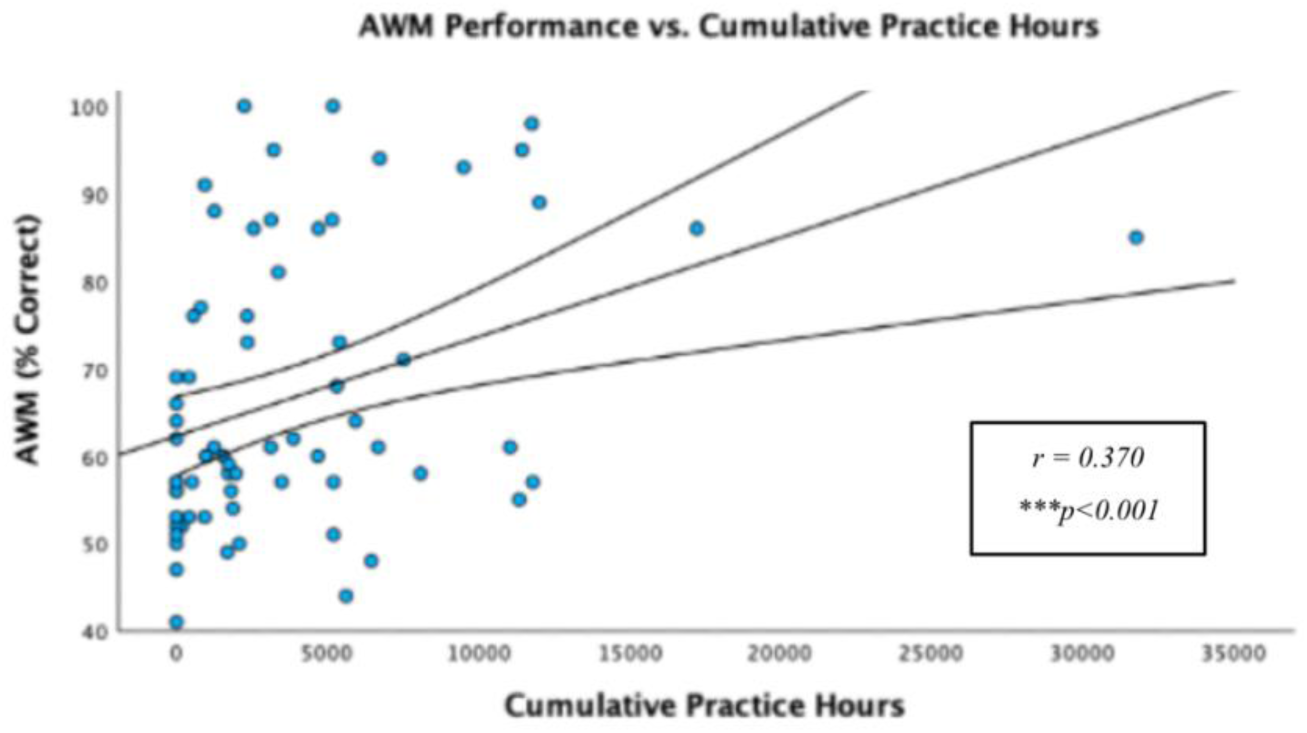
Experiment 2 Cumulative practice hours vs. AWM task performance. Pearson correlation is significant at the 0.01 level).

### Musical Training and MINT Outcomes

The mean accuracy score for the overall MINT was 73.73 (*SD* = 11.05). Independent samples t-test between Experiment 1 and 2 indicates a significantly lower MINT score for the subject in Experiment 2; t(144) = –5.36, p < .001. The mean accuracy scores for each MINT condition were: Baseline (Pitch) = 72.45 (*SD* = 16.25), Rhythm = 59.51 (*SD* = 12.95), Spatial = 71.76 (*SD* = 13.97), Visual = 83.43 (*SD* = 15.77), and Prediction = 81.47 (*SD* = 15.19). The mean accuracy scores for each SNR level were: SNR –3 = 79.29 (*SD* = 14.38), SNR –6 = 74.00 (*SD* = 12.51), and SNR –9 = 67.88 (*SD* = 13.04).

A correlation between cumulative practice hours and MINT task performance was tested with *r* = 0.293 (p = 0.008; **Figure 9**). Cumulative hours of practice were also correlated with the Baseline (Pitch) (*r* = 0.21, p = 0.040), Prediction (*r* = 0.29, p = 0.009), and Visual (*r* = 0.29, p < 0.009) sub-conditions. In addition, cumulative hours of practice correlated with SNR –3 (*r* = 0.34, p = 0.003) and SNR –9 (*r* = 0.21, p = 0.043). Independent samples t-test shows a significant difference in MINT performance between Musicians (mean = 78.74, *SD* =9.48) and Non-Musicians (mean = 66.15, *SD* = 11.54); t(35) = 3.63, p < 0 .001 (**Figure 10**).

**Figure 9.**
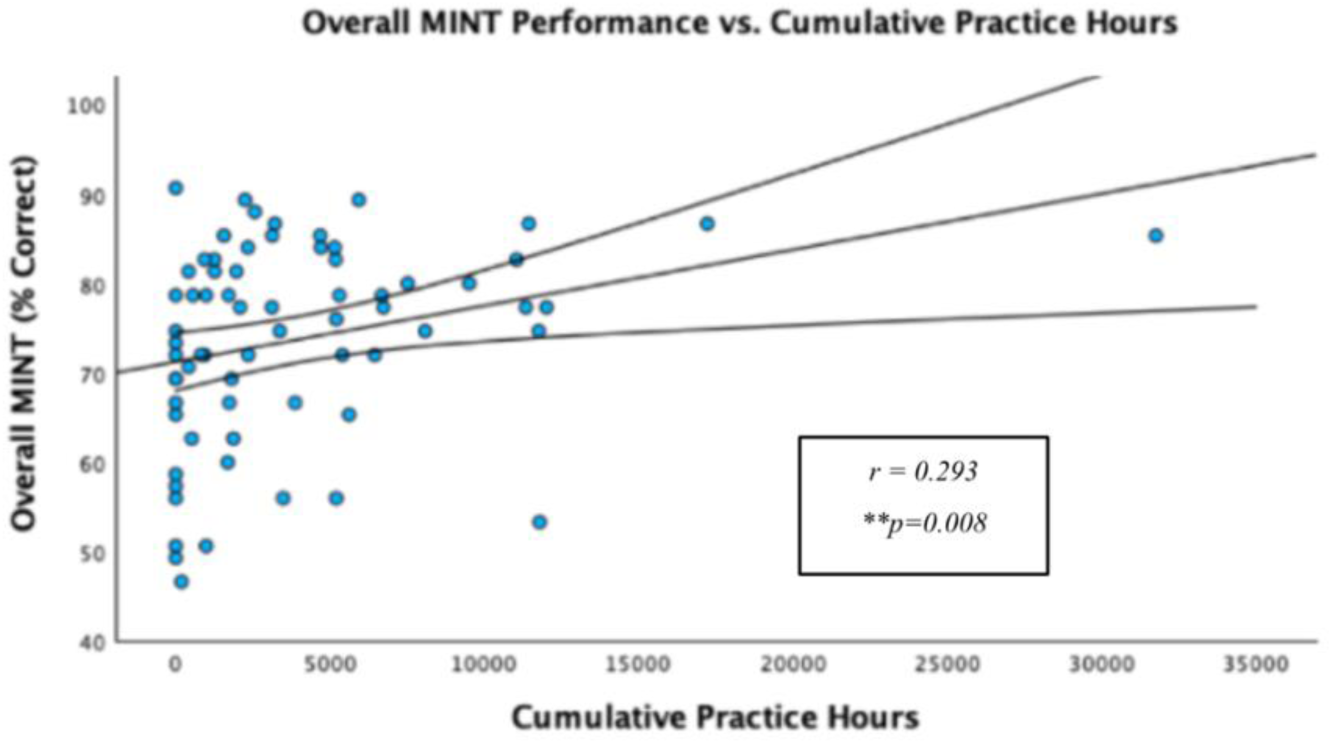
Experiment 2 cumulative practice hours vs. overall MINT performance. Pearson correlation is significant at the 0.01 level, N = 68.

**Figure 10.**
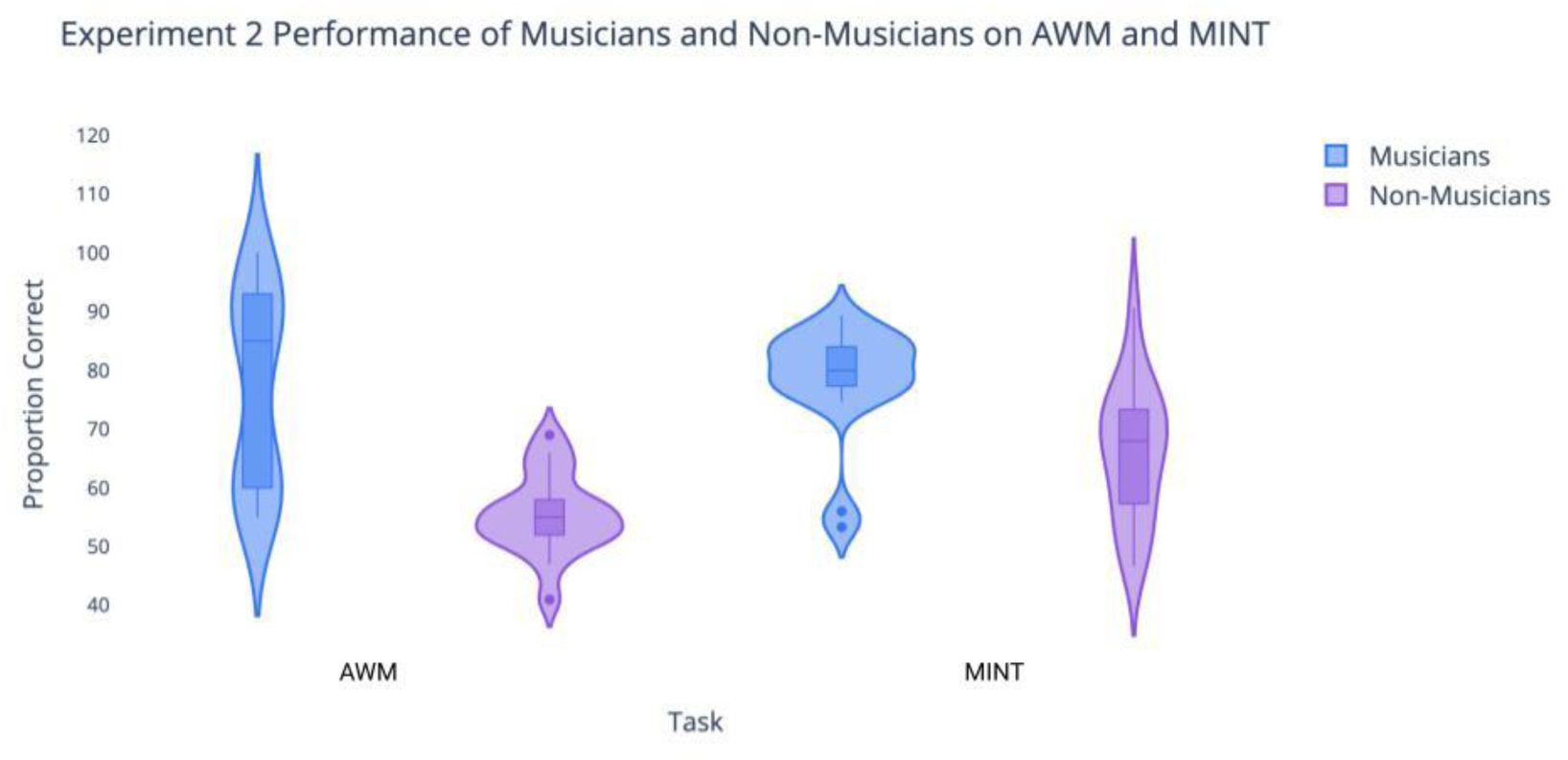
Violin plot showing Experiment 2 AWM task performance for Musician (mean = 76.68, *SD* =16.60, *N* = 19) and Non-Musician groups (mean = 55.44, *SD* = 6.82, *N* = 18). MINT performance for Musician (mean = 78.74, *SD* = 9.48) and Non-Musician groups (mean = 66.15, *SD* = 11.54). Significant group difference for both tasks *p* < 0.001.

### AWM and MINT Performance

Pearson correlation analysis evaluated the correlation between performance on the AWM and MINT tasks. AWM score significantly correlated with the overall MINT score (*r* = 0.573, p < 0.001; **Figure 11**). The correlations between AWM also significantly correlated with all MINT sub-conditions and SNR levels, as presented in **Table 3** and **Table 4**. Fisher’s test performed to compare the differences between the z-transformations of each pair of correlations demonstrated that none of the correlations are significantly larger than the others.

**Figure 11.**
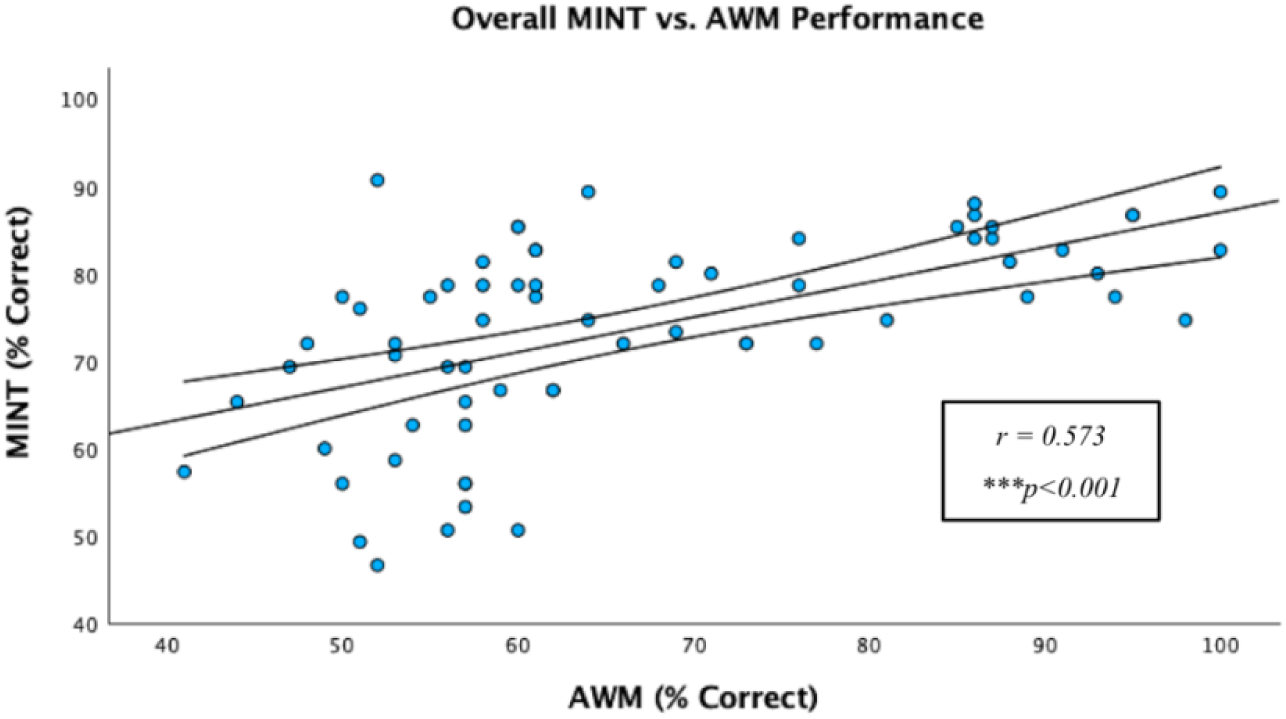
Experiment 2 AWM ability vs. overall MINT performance. Pearson correlation is significant at the 0.01 level, N = 68.

**Table 3.** Experiment 2 AWM Task Performance vs. MINT Conditions. Pearson correlations based on AWM task percent correct and average MINT scores for the corresponding condition.

|  | **Pitch**<br>**Total** | **Prediction**<br>**Total** | **Rhythm**<br>**Total** | **Spatial**<br>**Total** | **Visual**<br>**Total** | **MINT**<br>**Overall** |
| --- | --- | --- | --- | --- | --- | --- |
| **AWM Task Performance** | .451** | .424** | .292** | .438** | .507** | .573** |
| **Sig. (1-tailed)** | <.001 | <.001 | 0.008 | <.001 | <.001 | <.001 |

**Table 4.** Experiment 2 AWM Task Performance vs. MINT SNR Levels. Pearson correlations based on AWM task percent correct and average MINT scores for each SNR level across conditions.

|  | **SNR -9** | **SNR -6** | **SNR -3** |
| --- | --- | --- | --- |
| **AWM Task Performance** | .435** | .481** | .508** |
| **Sig. (1-tailed)** | <.001 | <.001 | <.001 |

### Mediating Role of AWM

Regression analyses were conducted to assess each component of the mediation model proposed in Experiment 1. Linear regression revealed a positive association between cumulative music training hours and both AWM performance (R = 0.370, *F*(1,66) = 10.48, p = 0.002) and MINT performance (R = 0.293, *F*(1,66) = 6.18, p = 0.015). AWM ability as the proposed mediator was also positively related to MINT test scores (R = 0.573, *F*(1,66) = 32.24, p < 0.001). Subsequent multiple regression analysis was performed to assess the effects of musical training hours (*X_1_*) and AWM (*X_2_*) on MINT performance (*Y*). The overall regression model was significant (R = 0.579, *F*(2,65) = 16.42, p < 0.001), with predictors contributing to improved MINT performance (β*_1_* = 0.093, p = 0.394; β*_2_* = 0.538, p < 0.001).

Given that the multiple regression model and the paths were significant and consistent with Experiment 1, mediation analysis was conducted using the same bootstrapping method with 5000 bootstrap samples. A 95% confidence interval for the indirect effect was derived from bootstrap samples and demonstrated a significant mediating role of AWM in the relationship between music training and MINT task performance. Results also show that the direct effect of music training on MINT became non-significant (p = 0.394) when controlling for AWM. Additional bootstrapping analysis also revealed a mediating effect of AWM on the Baseline (Pitch), Prediction, Spatial and Visual sub-conditions, and across all SNR levels.

## Discussion

### Effects of Musical Training on AWM and Music-in-Noise Perception

The findings from both Experiments 1 and 2 provide compelling evidence that supports our hypothesis of a musician’s advantage in both AWM abilities and music-in-noise perception. Importantly, the musician advantage was consistently observed across two distinct samples, which differed in overall musical experience, proportion of musicianship, and average performance on both tasks.

A meta-analysis conducted by Talamini et al. (2017) revealed that musicians outperform non-musicians in various memory domains, including long-term memory, short-term memory, and working memory. To examine AWM specifically, the task used in this study required participants to detect a local pitch change within two tonal patterns that differed in temporal order, assessing their ability to mentally manipulate the stimuli in addition to auditory retention (Albouy et al., 2017). Correlational analyses between cumulative practice hours and AWM task performance from Experiment 1 indicate a positive association between music experience and AWM abilities (**Figure 3**), a finding that is replicated in Experiment 2 (**Figure 8**). Moreover, the group comparison underscores this advantage, as musicians from both studies consistently outperform their non-musician counterparts on the standardized measures of AWM (**Figure 5**; **Figure 10**). These results are supported by existing literature, which consistently demonstrates behavioural, electrophysiological (event-related potential), and neuro-oscillatory evidence for the superiority of musicians in AWM abilities (Albouy et al., 2017; Foster et al., 2013; George et al., 2012;).

Analyses of the overall MINT task performance in Experiment 1 in relation to cumulative practice hours suggest a clear association between musical experience and improved music-in-noise perception (**Figure 4**). Although subjects in Experiment 2 showed an overall poorer performance in MINT—potentially due to differences in musicianship or increased task SNR difficulty—the correlation between musical experience and MINT task performance remained consistent and significant (**Figure 9**). In other words, the relationship between musical experience and music-in-noise perception is stable across different noise levels tested, irrespective of whether the SNR was set at 0, –3, –6 or –3, –6, –9. This finding illustrates that the influence of musicianship observed in the first experiment replicates despite heightened task demands and testing in a different sample. Furthermore, significant musician advantage on music-in-noise perception is also observed in both studies when comparing the musician and non-musician group differences in the overall MINT performance (**Figure 5**; **Figure 10**). These results are in line with the findings from Coffey et al.’s (2019) original MINT study and the subsequent MINT results by Hsieh et al. (2022), further validating the MINT task’s reliability and supporting the cognitive benefits of musical expertise amid varying levels of noise interference challenges.

### AWM Ability and Music-In-Noise Perception

Musicians’ music-in-noise benefits may arise from improvements in both auditory perception and cognitive processing. On the perceptual side, musicians demonstrate an increased sensitivity to fundamental acoustic features critical for music perception, such as pitch discrimination and temporal fine structure (Micheyl et al., 2006; Mishra et al., 2015). Cognitively, studies have shown a connection between musicianship and enhancements in cognitive faculties including working memory and attention (Bidelman & Yoo, 2020; Yoo & Bidelman, 2019), which may be linked to stream segregation improvements.

Evidence suggesting that AWM plays a crucial role in music-in-noise perception stems from the strong correlation between performance on the AWM task and the overall MINT score, observed in both Experiment 1 (**Figure 6**) and Experiment 2 (**Figure 11**). This finding replicates the original MINT study (Coffey et al., 2019), and is consistent with the majority of the SIN literature which suggests that working memory for phonological or tonal information is linked to improved speech segregation abilities (Bidelman & Yoo, 2020; Escobar et al., 2020; Lad et al., 2020; Mattys et al., 2012; Yoo & Bidelman, 2019).

The mediation analysis conducted in Experiment 1 further supports our hypothesis that AWM ability significantly mediates the relationship between musical experience and music-in-noise perception (**Figure 7**). This mediation model was successfully replicated in Experiment 2, with comparable results, reinforcing the reliability and generalizability of our initial findings. Overall, our results suggested that musicians’ enhanced AWM skills are a crucial driving force behind their enhanced MINT performance, and that musical training is associated with improvements in the performance of auditory stream segregation tasks through the enhancement of AWM capabilities. This mediating effect of AWM in music-in-noise performance parallels the mediation model proposed for AWM’s role in SIN performance (Kraus et al., 2012; Parbery-Clark et al., 2009b). Parbery-Clark et al. (2009b) demonstrated that musicians possess superior AWM skills, which they identify as a significant factor behind the group’s improved SIN performance. In addition, Bidelman & Yoo (2020) found that the relationship between musicianship and performance on a complex SIN task did not remain significant after controlling for working memory, which is associated with the listener’s year of musical training. This finding supports the concept that complex speech segregation is driven heavily by the enhanced working memory capacity, likely developed through musical training.

Although evidence supports the importance of AWM in overall stream segregation, the precise mechanisms underlying its contribution remain unclear. The predominant literature on SIN has focused on the role of AWM in facilitating the understanding of linguistic context (Kraus et al., 2012). For example, the Ease of Language Understanding (ELU) model by Rönnberg et al. (2013) posits that working memory enables the listener to hold a schematic representation of speech while processing contextual information, using linguistic knowledge to compensate for missing information in adverse listening environments. In addition, the ELU model states that individuals with enhanced working memory capacity spare more mental resources to resolve the phonological and semantic aspects of a listening task (Rönnberg et al., 2013). It follows that the advantage offered by AWM in aiding SIN processing may depend largely on the redundancy of linguistic contextual cues (e.g., phonological, lexical, syntactic, and semantic information) of the speech signal tested (Gordon-Salant & Cole 2016). However, given the consistent relationship between AWM and MINT performance —which is not influenced by linguistic factors—our study provides evidence that the benefits of AWM in stream segregation extend beyond the speech domain, pointing to more fundamental mechanisms that are universally involved in stream segregation processing.

### AWM’s Association with Perceptual and Cognitive Components in Stream Segregation

One advantage of the MINT is its ability to assess specific cues and auditory sub-skills related to stream segregation (Coffey et al., 2019), offering insights into how AWM may interact with the perceptual and cognitive elements involved in this process. The original MINT study indicated that AWM has the most significant contribution to the Prediction condition, and the relationship between musical training and the Prediction task diminished in significance when AWM performance was factored in as a covariate in the analysis (Coffey et al., 2019). Prior research also supports the role of AWM in predictive processing, highlighting its importance in top-down schematic expectations—the concept that anticipating the pattern to be segregated based on prior knowledge facilitates subsequent detection (Bey & McAdams, 2002).

However, contrary to earlier findings, we did not observe aa more important contribution of AWM to the Prediction condition compared to other conditions in our study. Instead, there were significant and consistent correlations between AWM and all MINT sub-tasks (**Table 1**; **Table 3**). This finding suggests that AWM’s contribution is not necessarily limited to conditions that specifically require retaining a melodic schema in working memory to anticipate incoming information, but rather represents a stable factor across various stream segregation situations.

One possible explanation for the contribution of AWM to general stream segregation is that enhanced AWM allows a more precise representation of acoustic signals in the mental workspace (Kraus et al., 2012). Research suggests that working memory is linked to improved performance on a rhythm synchronization task, where participants are required to reproduce the temporal structure of the presented rhythms (Bailey & Penhune, 2010). It is also indicated that individuals who can effectively retain auditory source properties, such as frequency and temporal fluctuations, over time have a perceptual advantage in SIN tasks (Lad et al., 2020; Lad et al., 2024). It is therefore plausible that the ability to maintain acoustic information accurately aids the sequential segregation processes essential for stream intelligibility (Bregman, 1990).

Another perspective involves attention. Dalton et al. (2009) manipulated the working memory load of participants while they performed a distractor interference task, demonstrating a causal role for the availability of working memory in auditory selective attention. In addition, it is suggested that segregating auditory streams from background noise draws upon attentional resources (Heinrich et al., 2008), and accomplishing such tasks necessitates the allocation of one’s limited cognitive resources to balance the competing demands of attention, processing, and storage (Wingfield & Tun, 2007). It is therefore plausible that enhanced AWM proficiency promotes the maintenance and encoding of auditory signals, which in turn allows for more efficient use of attention resources to extract and recall the target stream.

Alternatively, the advantages of AWM can be understood through the concept of temporal integration. It is proposed that working memory aids the logical and behavioural linkage between recent past and imminent future events, thus serving both a retrospective role in information retention and a prospective role in anticipation (Fuster & Bressler, 2012).

Specifically, prior literature proposes that the availability of working memory is important for minimizing distractor interference through the active maintenance of current stimulus-processing priorities (Dalton et al., 2009; Lavie, 2005). In stream segregation, AWM may therefore enable individuals to hold fragments of auditory information while processing, integrating, and anticipating degraded target signals. Future research is suggested to investigate the nuanced mechanisms of AWM in aiding the segregation of an intended auditory signal in challenging acoustic situations.

### The Auditory Dorsal Stream and its Implications for Musician Enhancement

The dorsal stream of auditory processing, which involves the parietal lobe, dorsal premotor cortex, and dorsolateral frontal regions, is central to higher-order cognitive auditory functions. It supports the manipulation of sound patterns in working memory, auditory-motor integration, abstract temporal representations, and predictive coding (Rauschecker & Scott, 2009; Zatorre, 2024). Neuroimaging studies highlight the dorsal stream’s key role in AWM, with activations in parietal regions associated with mental transformation and manipulation processes (Foster et al., 2013; Zatorre et al., 2010). For instance, an fMRI study shows that the intraparietal sulcus (IPS) within the dorsal stream is engaged during pitch and temporal transformations of auditory representations (Foster et al., 2013). Moreover, Albouy et al. (2017) have observed that sustained evoked activity in the bilateral dorsal streams, particularly through long-range theta phase locking and increased local theta power in the IPS, is associated with successful AWM manipulation. Furthermore, when theta power is boosted in the dorsal stream via rhythmic brain stimulation (Albouy et al 2017) or via flickering visual stimulation (Albouy et al 2022), AWM performance is also enhanced.

While perceiving auditory signals in background noise heavily engages primary and non-primary auditory regions (Holmes et al., 2021; Kell & McDermott, 2019), research indicates that motor and somatosensory areas are also more actively recruited under challenging listening conditions (for review, see Skipper et al., 2017). This suggests a compensatory mechanism of dorsal steam activity for reduced processing specificity in the auditory system (Du et al., 2014). Importantly, a study comparing musicians and non-musicians found that the benefits of musical training on SIN perception in difficult listening contexts were related to activity in the motor cortices of the auditory dorsal streams (Du & Zatorre, 2017).

Further research has shown that music training enhances functional connectivity within the dorsal auditory stream (Jünemann et al., 2013). Musicians also exhibit greater structural connectivity in the white matter tracts of the dorsal stream (i.e., arcuate fasciculus and superior longitudinal fasciculus; Halwani et al., 2011; Oechslin et al., 2010). Differences in the microstructural plasticity of dorsal white matter are suggested to underlie musicians’ improved SIN perception (Li et al., 2021). Considering the role of the auditory dorsal stream in AWM and SIN perception, we thus infer that the musician enhancements in these abilities may be rooted in this stream, although the exact mechanisms warrant further exploration.

### Implications for Age-Related Hearing Loss

Auditory functioning is one of the most prevalently affected sensory modalities in the elderly population (Yamasoba et al., 2013), with age-related hearing loss impacting over 5% of the global population (Davis et al., 2016). In addition, older adults, with or without hearing loss, show disproportionate deficits in speech recognition in noisy environments and AWM compared to younger adults (Dubno et al., 1984; Humes & Floyd 2005). Previous studies showed that older musicians exhibit enhanced performance in AWM and SIN perception compared to their non-musician counterparts, suggesting that musical experience may mitigate age-related hearing challenges (Zhang et al., 2021; Zendel et al. 2019).

Recent longitudinal studies assigning older adults to musical activities (piano/choir) have also demonstrated behavioural, neurophysiological, and neuro-oscillatory evidence of improvements in SIN perception (Worschech et al., 2021; Hennessy et al., 2021; Dubinsky et al., 2019; Gray et al., 2022). Shedding light onto the mediating role of AWM in stream segregation, we propose that future music programs designed to address hearing challenges in older adults should focus on enhancing AWM to achieve optimal intervention outcomes.

### Limitations and Future Directions

Limitations of the current study include reliance on self-report music history questionnaire responses and the challenge of precisely controlling for the nuanced variations of individual musical experiences and expertise (e.g., learning styles, extent of practice). Moreover, the correlational design of the study does not address issues regarding self-selection and the direction of causality, particularly considering evidence suggesting that auditory and musical expertise arises from a combination of genetic predispositions and experiential plasticity (Schellenberg, 2015; Zatorre, 2013). The inherent predispositions for AWM or stream segregation abilities could potentially influence one’s path toward musical engagement, an aspect that warrants further investigation.

Longitudinal studies with school-aged children (as well as the elderly, as described in the preceding section) provide evidence that music instruction is in fact causally associated with moderate benefits in SIN and AWM abilities (Slater et al., 2015; Nie et al., 2022), but of course, this does not mean that predisposing factors do not exist. Moreover, while a correlation between MINT and SIN performance has been observed (Coffey et al., 2019), suggesting some shared contributions, the extent of transfer effects from music to speech remains unclear. Therefore, future research directions entail conducting longitudinal studies to examine the development of both speech-in-noise and music-in-noise perception, further unravelling the relationship between musical training, AWM, and overall auditory stream segregation. Such endeavours will also help elucidate experience-dependent plasticity in the auditory domain and contribute to a deeper understanding of the development of higher-level auditory cognitive mechanisms.

### Conclusion

This study explores the influence of musical training on two auditory cognitive processes: AWM and stream segregation. As hypothesized, our findings provide support for a musician advantage in AWM abilities and music-in-noise perception. We show using replication across two samples that musicians’ enhanced AWM skill is one of the driving forces behind their better music-in-noise performance, suggesting that musicianship fosters improvements in stream segregation through the enhancement of AWM capabilities. In addition, the study’s two-phase design strengthens the generalizability of the results across various populations and conditions. Together, these findings shed light on the relationship between musical training, AWM, and stream segregation, underscoring the potential for music-based interventions to enhance auditory processing abilities.

## Acknowledgments

This research was supported in part by funding from the Canada First Research Excellence Fund, awarded to RJZ and JG via the Healthy Brains, Healthy Lives initiative at McGill University and BrainsCAN at Western University, and also by a grant from the Canadian Institutes for Health Research and the Grand Prix Scientifique from the Fondation pour l’Audition to RJZ. The authors thank Emmett Lewis-Hoeber, Sebastian Kolde, Ethan Yan, Amy Li, Lucy Core, and Emily Chen for their assistance in data collection, and Philippe Albouy for providing the task stimuli.

